# Differential response trajectories to acute exercise in blood and muscle

**DOI:** 10.1101/863100

**Authors:** David Amar, Malene E. Lindholm, Jessica Norrbom, Matthew T. Wheeler, Manuel A. Rivas, Euan A. Ashley

## Abstract

A physically active lifestyle is essential for maintaining health, and is a powerful way to prevent chronic disease. However, the molecular mechanisms that drive exercise adaptation and transduce its beneficial effects, are incompletely understood. Here, we combined data from 49 studies that measured the whole transcriptome in humans before and after exercise to provide the power to draw novel observations not seen in any individual study alone. The resulting curated and standardized resource includes samples from skeletal muscle (n=1,260) and blood (n=726) in response to endurance or resistance exercise and training. Using a linear mixed effects meta-regression model selection strategy, we detected specific time patterns and novel regulatory modulators of the acute exercise response. Acute and long term responses to exercise were transcriptionally distinct. Exercise induced a more pronounced inflammatory response in skeletal muscle of older individuals. We identified multiple sex-specific response genes, where *MTMR3* is a novel exercise-regulated gene. These results deepen our understanding of the transcriptional responses to exercise and provide a powerful resource for future research efforts in exercise physiology and medicine.

## Introduction

A physically active lifestyle is one of the most important actions by which individuals of all ages can improve their health. It prevents common chronic diseases, reduces anxiety, and improves cognitive functions and the overall quality of life (Neufer et al., 2015; Piercy et al., 2018). Both acute effects and long-term adaptation have been studied in response to endurance (repeated contractions of low force) and resistance (fewer contractions of greater force) exercise in humans, where skeletal muscle and blood are the most accessible and well-studied tissues.

Exercise-induced adaptation occurs in working skeletal muscles, however multiple myokines are released from skeletal muscle into blood, with the potential to affect all organs (Pedersen and Febbraio, 2012). Resistance training specifically leads to skeletal muscle hypertrophy, while endurance training leads to increased capillary density and ATP-producing capacity. Both exercise modalities improve insulin sensitivity (Di Meo et al., 2017), decrease blood pressure (Cornelissen and Smart, 2013), and improve the blood lipid profile by increasing HDL cholesterol and reducing LDL and triglyceride levels (Cornelissen et al., 2011; Mann et al., 2014; Seals et al., 1984).

Exercise-induced change in gene expression is an important aspect of adaptation. The global transcriptional response to both acute exercise and long-term training, measured using microarrays or RNA-sequencing, has been investigated in healthy humans by sampling human blood (Nakamura et al., 2010; Neubauer et al., 2013; Radom-Aizik et al., 2008; Rampersaud et al., 2013) and skeletal muscle (Huffman et al., 2014; Keller et al., 2011; Lindholm et al., 2014a; Raue et al., 2012; Robinson et al., 2017; Vissing and Schjerling, 2014). These studies identified multiple differentially expressed genes. Although resistance and endurance training lead to distinct cellular adaptation, there are several genes, including known regulatory factors, that are differentially expressed in response to both modalities, for example *VEGFA*, *PDK4*, *PPARGC1A* and *FOXO3* (Barrès et al., 2012; Lindholm et al., 2014a; Raue et al., 2012). Similar findings were reported by Lundberg et al. (Lundberg et al., 2016) on the global transcriptional response to combined resistance/endurance training.

All studies above are limited in size due to the complexity of conducting controlled interventions in healthy individuals that include invasive biological sampling. Consequently, available datasets differ substantially in the clinical attributes of their subject sets, including: sex, age, training modality, and sampled post-exercise time points. Thus, each study alone is limited in the ability to reveal genes that are consistently differential and are associated with these variables.

Meta-analysis is a standard tool for systematic quantitative analysis of previous research studies. It can generate precise estimates of effect sizes and can be more powerful than any individual study contributing to the pooled analysis (Haidich, 2010). Moreover, the examination of variability or heterogeneity in study results is also critical, especially when apparently conflicting results appear. Meta-analysis can be based on simple weighted averaging of effects using random effects (RE) models, or, in more complex cases, be based on meta-regression of effect sizes vs. covariates, which are typically called *moderators*. Meta-analysis (and meta-regression) has been traditionally used in medicine and epidemiology, but it has been gaining popularity in large-scale genomics studies (Ramasamy et al., 2008; Sweeney et al., 2017).

Here, we gathered and annotated publicly-available transcriptome datasets from both endurance and resistance exercise interventions in humans. This allowed us to access a larger number of samples (1986; 726 from blood and 1260 from skeletal muscle) compared to previously published studies of the exercise transcriptome on 120 samples in blood (Rampersaud et al., 2013) and 119 samples in skeletal muscle (Lindholm et al., 2016). Our results expand the understanding of the transcriptional landscape of exercise adaptation by extending previously known expression responses and their regulatory networks, and identifying modality-, time-, age-, and sex-associated changes. For time-associated gene modules, we detect novel regulators of the acute exercise response.

## Results

### Data collection and analysis

We collected and annotated data from 49 human exercise training studies, including 2451 samples from 1450 subjects in total. See Figure 1 for an overview of the study. Dataset records from the Gene Expression Omnibus (GEO), manuscripts, supplementary information, and personal communication with authors were used to extract sample-level or study-level information including sex, age, tissue, training modality, additional treatments, and time points. Most of the data (1986 samples from 959 subjects) were from studies that had both pre- and post-exercise data and sampled blood and/or muscle. Transcriptome data from these 1986 samples were used for the meta-analysis. Of the 959 subjects, 142 did not have sex information and 397 of the remaining (47%) were female. For samples with missing sex annotation, biological sex was imputed using expression of Y chromosome genes (internal validation was at 0.99 ROC, see **Materials and Methods**). Each dataset was partitioned into homogeneous cohorts that had a similar training regime, see Supplementary Table 1 for details. This resulted in 81 cohorts, only two of which were associated with a disease (COPD, a subset of study GSE27536) or a post-exercise sample treatment (endotoxin, a subset of study GSE83578) and were included in the meta-analysis. For each gene and a post-exercise time point, in each cohort, we computed the effect size of differential expression compared to the pre-exercise sample. In addition, for each cohort we kept the following four *moderators* (covariates): average age, proportion of males, time points, and training modality.

**Figure 1.**
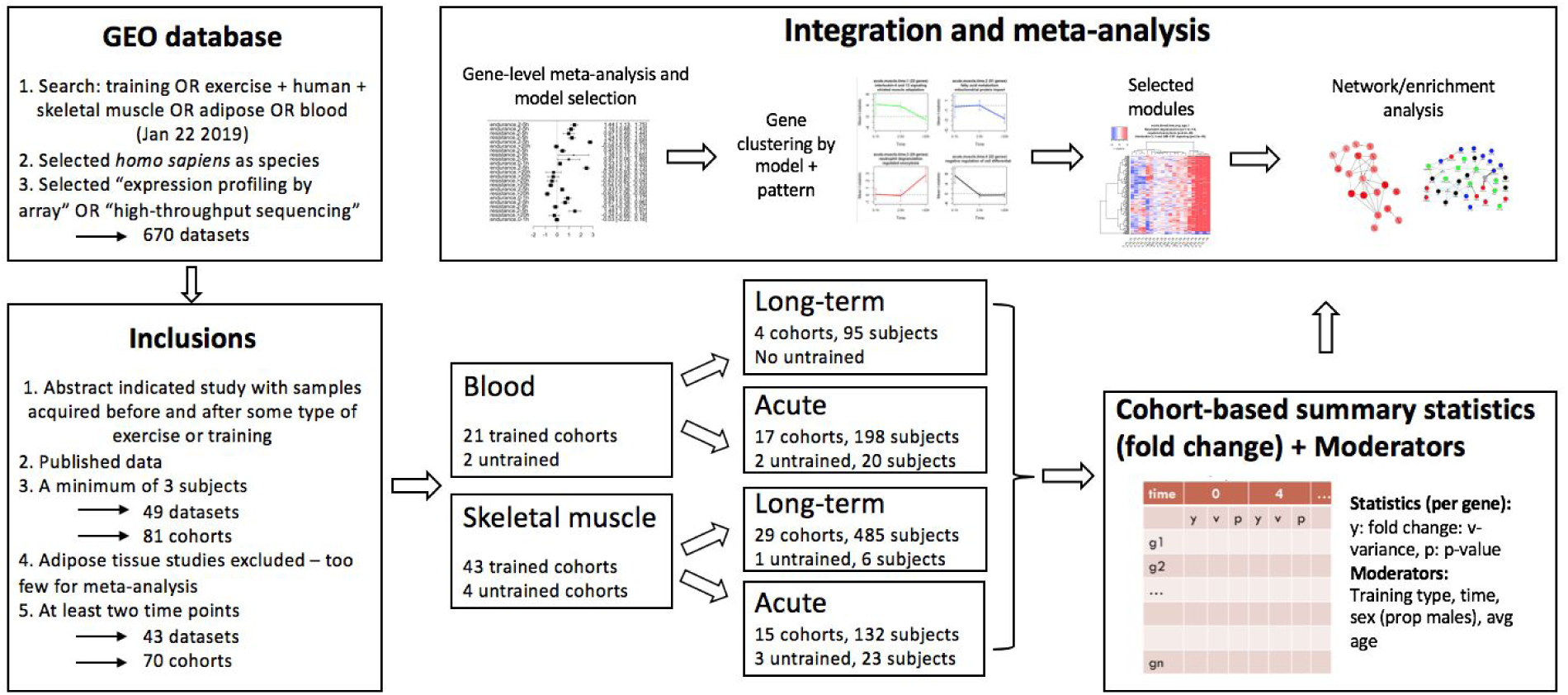
Study overview. We started with a search in the Gene Expression Omnibus. Manual examination of the studies and a set of filters resulted in 43 studies that had whole-genome expression profiles from blood and muscle. The data covered 70 exercise cohorts that were partitioned into four types of meta-analysis (i.e., by tissue and exercise modality). Summary statistics and moderators were computed for each cohort and were used as input for the statistical analysis. Gene-level model selection and meta-analysis was used to detect differential genes, which were later clustered by their patterns. Network and enrichment analysis were used for interpretation.

To keep the input for the meta-analyses homogeneous, we partitioned the analysis into four types by tissue (blood or muscle) and intervention (acute exercise bout vs. long-term training). Figure 2A illustrates the high statistical heterogeneity of the resulting datasets when performing a simple random effects (RE) meta-analysis for each gene. This high heterogeneity likely resulted from having both high clinical and statistical heterogeneity. The clinical heterogeneity stems from both having studies that were conducted in different populations and unbalanced sampling of moderators. For example, acute bout skeletal muscle studies were heavily biased towards men, and time coverage was not balanced across the acute studies, see Supplementary Table 1. We also examined the associations between the moderators across the cohorts, see Supplementary Figure 1. Generally, except for high bias in time window 0-1h towards young individuals in acute blood cohorts, most associations were mild (median R^2^ < 12%) with only a few that were significant.

**Figure 2.**
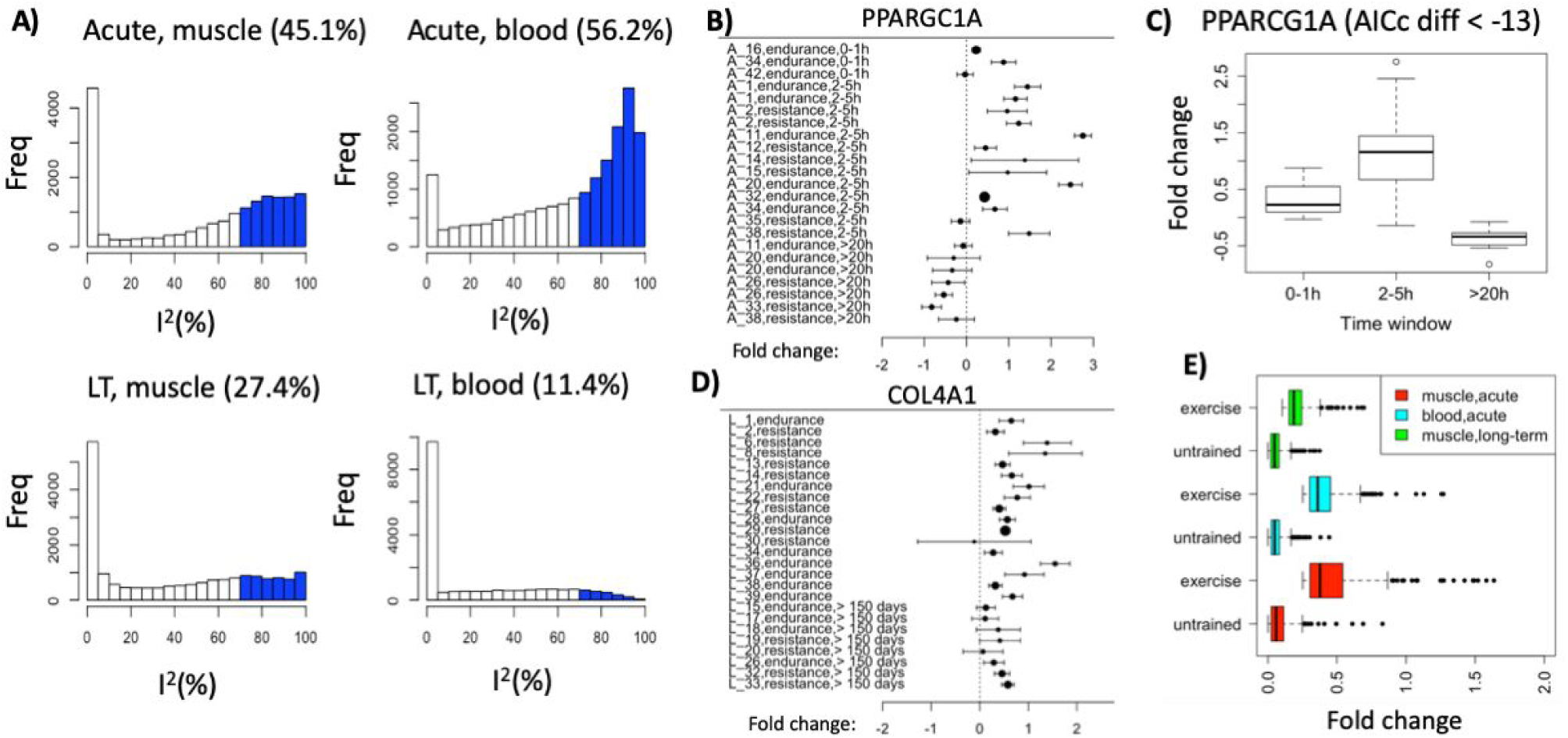
Properties of the meta-analysis results. A) Most genes have high true heterogeneity with high heterogeneity of true effects (I^2^>70%) bins marked in blue (the percent of these genes appears in the title of each plot). LT: long-term training. B) Forest plot of the effect sizes of *PPARGC1A* in acute muscle cohorts. C) Illustration of the fold change across time. Boxplots summarize the fold changes over the cohorts of the time window. D) Forest plot of the effect sizes of *COL4A1* in long-term training studies in skeletal muscle. Upregulation is consistent across the cohorts and the random effects estimation is presented below the plot (RE model). E) The differential expression effects of our selected genes is unique to exercise. Colors represent the different meta-analysis types in the study. For each meta-analysis the effect sizes of our selected genes is presented, once for the exercise cohorts and once for the untrained cohorts.

Supplementary Figure 2 illustrates the inflation of low p-values of the random effects meta-analyses, indicating a high false positive rate if RE models were to be used alone. Such inflation is expected when the heterogeneity of true effects is high (as illustrated in Figure 2A).

To mitigate these issues we developed a pipeline that fits a model for each gene (see **Materials and Methods** for details). Briefly, for each gene we considered different combinations of the moderators, each producing a different meta-regression model. The model without any moderators was fit using RE meta-analysis and is denoted as the *base model*. Model selection was performed using the Akaike information criterion with correction for small sample sizes (AICc) (Burnham and Anderson, 2002; Cavanaugh, 1997). However, to focus on robust genes, we applied an additional series of conservative filters, including both a high inferred fold change (>0.25 for acute and >0.1 for longterm) and a high significance of the top model (p<1×10^-4^, corresponds to FDR<0.1 over all tests). Moreover, a model was considered a significant improvement over the base model only if the AICc improvement was >5 (Supplementary Figure 3). Otherwise, we took the base model, but only if the inferred variance was reasonably low (I^2^ < 50%).

To illustrate the output of the model selection process, consider the response of *PPARGC1A* (PGC-1ɑ) in skeletal muscle after acute exercise. This gene is an exercise-mediated regulator of muscle adaptation with a well-documented transcriptomic response (Lira et al., 2010). In our analysis, *PPARGC1A* passed all statistical tests above and its top model included time as a moderator. The forest plot (Figure 2B) shows a consistent upregulation at the intermediate (2-5h) time point following exercise, with no change immediately after exercise and a small downregulation at later time points (>20h), as summarized in Figure 2C. As another example, consider *COL4A1*, a type IV collagen gene, which showed a consistent induction with long-term training across studies (Figure 2D).

The model selection process detected differential genes in each of the four analyses: acute exercise, muscle: 223 genes; long-term training, muscle: 334 genes; acute exercise, blood: 388 genes; and long-term training, blood: seven genes. Most detected genes had one or more moderators in their selected model. Supplementary Figure 4 shows the statistics of the selected models and their moderators (see Supplementary Table 2-6 for full details). Results for long-term blood studies are not shown as only base models were selected (see Supplementary Table 6). The results show which moderators were selected together in the models of the genes. For example, four genes had models that had both time and training features in acute skeletal muscle studies (Supplementary Figure 4A). Note that many co-occurring moderators were related to time. Specifically, our meta-regression tested for both linear and quadratic trends. For many models, both of these patterns were selected. An example of a gene with such a model is shown in Supplementary Figure 4D. Here, linear or quadratic patterns alone cannot fully capture the time-associated pattern in which the gene is upregulated at the 0-1h window but has no differential expression in the next time windows.

In order to validate the results of our model selection pipeline we performed two additional analyses. First, we performed a meta-analysis of the untrained cohorts, which were ignored in the main analysis above. We compared the inferred effect sizes of our selected genes above (i.e., 223 from the acute muscle analysis, 334 from long-term muscle, and 388 from acute blood) in the exercise cohorts with the untrained cohorts, see Figure 2E. The results show that the exercise effects are substantially greater than the effects in the untrained controls (all paired tests were significant at p<1×10^-05^). Second, we quantified the replicability of the genes based on their p-values across the time points and cohorts. We used the SCREEN algorithm (Amar et al., 2017) to estimate replication across the studies, while modeling the latent correlation structure of the cohorts, see **Materials and Methods** for details. Supplementary Figure 5 shows the local FDR scores of our gene set as compared to all other genes, which are consistently much lower. This analysis illustrates that our gene sets perform well both in FDR control and in replication across four or more cohorts.

### Co-expressed gene clusters reveal exercise response pathways

The results above were based on the data of a single gene at a time. To allow for a more holistic view of the biological processes that govern the transcriptomic response to exercise, we clustered the selected genes by their differential expression patterns. We first grouped the genes by their selected model types. For example, we had a set of 123 genes in the acute muscle meta-analysis with only time as the selected moderator. We then clustered these gene sets using the K-means algorithm, applied to their average summary statistics across the cohorts, see **Materials and Methods**. The 123 genes clustered into four groups based on their time courses (Figure 3A). In total, 17 out of 26 detected gene clusters with at least 5 genes had at least one significantly enriched GO term or Reactome pathway (10% FDR adjustment), see **Supplementary Tables 7-8**, illustrating their high quality and agreement with known biological function.

**Figure 3.**
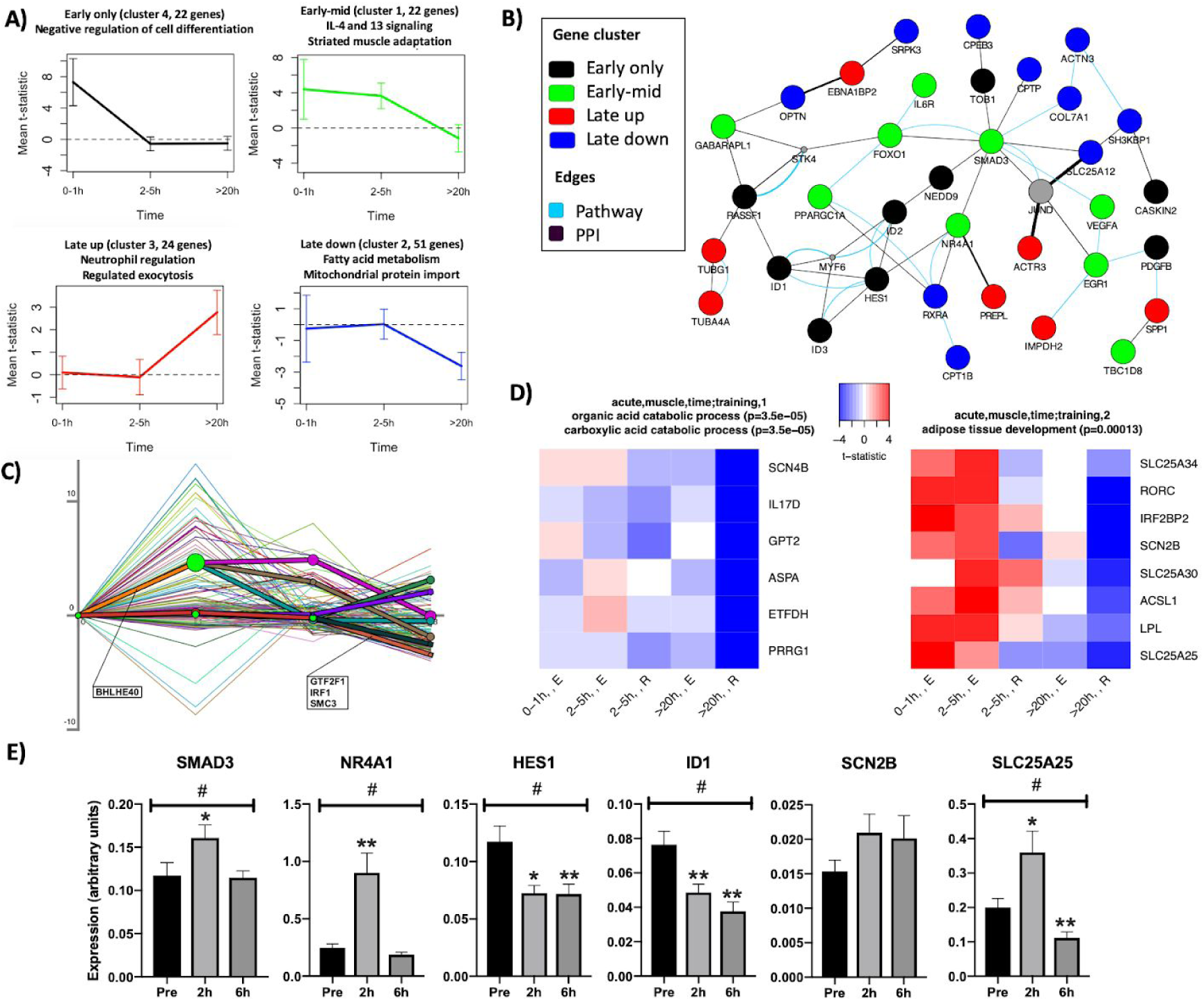
Differential expression patterns in skeletal muscle after acute exercise. A) Genes associated with time were partitioned into four groups based on their trajectories. The title of each subplot shows significantly enriched GO-terms or pathways. B) The main GeneMANIA connected component of the genes in A when overlaid on known protein-protein or pathway networks. Gray nodes are genes predicted to be functionally relevant because of their high connectivity. C) DREM analysis results of the genes in A predict several transcription factors to be involved in the observed responses. D) Gene modules associated with both time and training modality (endurance - E, resistance - R). E) Validation of gene expression changes following acute endurance exercise in a separate human cohort. SMAD3, NR4A1, HES1 and ID1 from the acute network and SCN2B and SLC25A25 from the endurance and time-specific genes. # denotes an overall significant treatment effect between time points (*P*<0.005). *(*P*<0.05) and **(*P*<0.005) denote significant expression difference compared to before exercise (pre) after correction for multiple comparisons. Values are presented as mean±SE.

### Temporal patterns imply information propagation across the gene network of muscle response to acute bouts

As explained above, the 123 genes associated with time in the acute skeletal muscle analysis were partitioned into four clusters, a novel representation of exercise-induced gene expression changes where each cluster corresponds to a different time trajectory and a different functional annotation, see Figure 3A. To better understand the functional implications of these, we also overlaid the genes on a network of known pathway or protein-protein interactions (PPI), see Figure 3B. This resulted in a large connected component with representation from all four clusters. Genes from the clusters with a late up- or downregulation response had low degrees (at most 3 neighbors, genes from clusters 1 and 3), whereas genes from the early response clusters formed hubs and were well connected. This network structure suggests that the main hubs respond first and their effect is then propagated downstream.

We used DREM (Schulz et al., 2012) as a complementary approach to cluster temporal responses. DREM integrates the time courses with known transcription factor-target networks and sequence information to produce a model that links temporal responses to putative regulators. In our data, DREM predicted BHLHE40 to be an important upstream transcriptional regulator during the early 0-1h time window (Figure 3C), a novel observation in this study. *BHLHE40* is itself induced by a bout of combined aerobic and resistance exercise in human skeletal muscle (Lundberg et al., 2016). *BHLHE40* is induced by hypoxia and directly binds to PGC-1ɑ and represses its transactivation activity, likely to reduce hypoxia-induced ROS damage. The repression of PGC-1ɑ by BHLHE40 has been shown to be alleviated by an increase in *PPARGC1A* expression itself *in vitro*, and by running exercise in mice (Chung et al., 2015). DREM analysis also predicted three different transcription factors to be involved in the regulation of the late response genes (Figure 3C). The transcription factor IRF1 (Interferon Regulator 1) has been identified previously as an upstream regulator of the transcriptional endurance training response (Keller et al., 2011), which corroborates the DREM predictions. Another predicted regulator was GTF2F1, which is a general transcription factor that for example activates serum response factor (Joliot et al., 1995), that is activated by exercise (Rose et al., 2006).

The temporal cluster analysis identified key genes known to be regulated by acute exercise in humans. The early only response gene cluster was functionally enriched for negative regulation of cell differentiation, which includes a significant induction of the LDL receptor (*LDLR*) gene. An immediate change in *LDLR* expression in muscle has been demonstrated after a 45 min cycling bout (Pourteymour et al., 2017) and with 3 months of endurance training (Lindholm et al., 2014a). An increase in LDL receptor abundance in skeletal muscle could lower plasma LDL cholesterol, which is essential for preventing cardiovascular disease (Study of the Effectiveness of Additional Reductions in Cholesterol and Homocysteine (SEARCH) Collaborative Group et al., 2010), and is a well-established effect of exercise training (Albarrati et al., 2018). Genes related to angiogenesis were also induced early, for example *PDGFB* (Platelet-Derived Growth Factor B) and *VEGFA* (Vascular Endothelial Growth Factor A) that belong to the same protein family. Both factors also increase in blood after strenuous exercise (Czarkowska-Paczek et al., 2006). Another early-response growth factor was *EGR1* (Early Growth Response 1).

Transcriptional regulation associated with striated muscle adaptation occurred during the 0-5h time period following acute exercise (early-mid response cluster). PGC-1ɑ, a central regulator of mitochondrial biogenesis, belongs to this group. PGC-1ɑ is a transcriptional co-regulator that is well-studied in relation to exercise adaptation (Egan and Zierath, 2013; Lira et al., 2010). It acts through co-activation of several key transcription factors including PPARs for induction of *GLUT4* for glucose uptake, NRF1 for mitochondrial biogenesis through *e.g. TFAM* and cytochrome C activation (Herzig et al., 2000; Virbasius and Scarpulla, 1994), and ERRɑ for activation of *VEGF* transcription, which promotes angiogenesis (Arany et al., 2008). *FOXO1* had a similar induction pattern. It interacts with PGC-1ɑ to induce expression of lipid metabolism genes in response to exercise-induced mechanical stretch. One of the central hubs of the acute response network is *SMAD3*, an intracellular effector of TGF-β. *SMAD3* is required for the atrophic effect of Myostatin/TGF-β (Sartori et al., 2009), and both FOXO1 and SMAD3 regulate myostatin expression (Allen and Unterman, 2007). *SMAD3* has shown a greater induction in response to resistance training in females compared to males (Liu et al., 2010). These central factors regulate the balance between protein synthesis and degradation, which is key for the hypertrophic response to resistance training.

*SPP1* is an example of a gene that increased in the late time window (>20h post exercise) (Figure 3A and B). Its expression is stimulated by PGC-1ɑ, and the SPP1 protein is subsequently secreted and has been shown to activate macrophages to induce angiogenesis in murine skeletal muscle (Rowe et al., 2014). Genes related to fatty-acid metabolism were downregulated during the later time points following acute exercise (Figure 3A, late down cluster). Examples of genes with this expression pattern are *ACOT1*, an Acyl-CoA thioesterase; *MLYCD*, which produces Acetyl-CoA from Malonyl-CoA and thus stimulates fatty acid oxidation; and *CPT1B*, which is a mitochondrial membrane transporter of fatty acids.

The genes discussed above were discovered in clusters associated with time only. However, we detected two additional gene modules associated with time and training modality, see Figure 3D. The first module (Figure 3D, right panel) depicts an early endurance-specific upregulation response enriched with fatty acid metabolism functions. Specifically, lipoprotein lipase (LPL) is a key enzyme for the breakdown of circulating triglycerides for uptake of FFAs into skeletal muscles and ACSL1 has been shown in mice to be essential for preserving blood glucose levels during exercise through stimulation of beta-oxidation (Li et al., 2015). Several mitochondrial transporters also showed differential regulation specific to training modality, e.g. *SLC25A25* that is important for ATP homeostasis. The left panel of Figure 3D shows a downregulation of genes associated with catabolic processes specifically in response to resistance exercise. A shift in balance that favors anabolic processes is known to occur with hypertrophy-inducing resistance exercise (Damas et al., 2015).

We used a separate human acute exercise cohort to validate the findings from the meta-analysis with qRT-PCR. Sixteen subjects, 8 males and 8 females, performed 1h of endurance exercise at 70% of their peak VO_2_. Skeletal muscle biopsies were obtained before, and at 2 hours and 6 hours after exercise. SMAD3 and NR4A1 were validated as part of the central acute cluster network, and in accordance with the meta-analysis findings, both were significantly upregulated at the 2h time point following exercise. HES1 and ID1 were induced in the early 0-1h window in the meta-analysis, which was not covered in the validation cohort. However, both genes showed significant positive quadratic, and negative linear trends in the meta-analysis. Supplementary Figure 6 illustrates how these genes are mostly down-regulated in the 2-6h window. Our validation showed a significant downregulation at both 2h and 6h after exercise, which is in agreement with the meta-analysis. SCN2B and SLC25A25 were selected as endurance-specific genes. Both showed significant upregulation in the early time windows following exercise. No significant upregulation was observed for SCN2B, although there was a trend towards an increase. SLC25A25 increased at the 2h timepoint, while a decrease was observed at 6h, in concordance with the >20h time window for the meta-analysis.

### Long-term training induces extracellular matrix reorganization genes in skeletal muscle

The long-term training studies included in the meta-analysis ranged from 6 weeks to 9 months of training. Exercise-modality independent response analysis identified six downregulated genes (Figure 4A) and 60 upregulated genes (Figure 4B) associated with long-term training in skeletal muscle (Supplementary Table 5). Myostatin (*MSTN*), which is well-characterized as a negative regulator of skeletal muscle hypertrophy, was one of the downregulated genes. Downregulation was also seen for *BCL6*, a germinal center B cell and follicular helper T cell transcriptional regulator, which is consistent with data showing inverse correlation between its expression levels and VO_2_max in humans (Parikh et al., 2008). The upregulated genes were enriched for extracellular matrix (ECM) reorganization (Figure 4B), a central training adaptation mechanism (Keller et al., 2011; Martinez-Huenchullan et al., 2017). Several collagen genes important for ECM, including *COL4A1*, *COL4A2*, *COL1A1* and *COL6A3* were induced. *COL4A1* represents one of the major components of the basement membrane surrounding skeletal muscle fibers (Gillies and Lieber, 2011). In addition to endurance and resistance training, *COL4A1* expression is also induced by high-intensity interval training (Miyamoto-Mikami et al., 2018), and its expression response has been associated with high- and low-responders to endurance training (Keller et al., 2011). Based on the meta-analysis findings, we can expand from the current knowledge of this network as displayed in Figure 4C to an improved, more comprehensive undirected interactome for long-term transcriptional response to training (Figure 4D).

**Figure 4.**
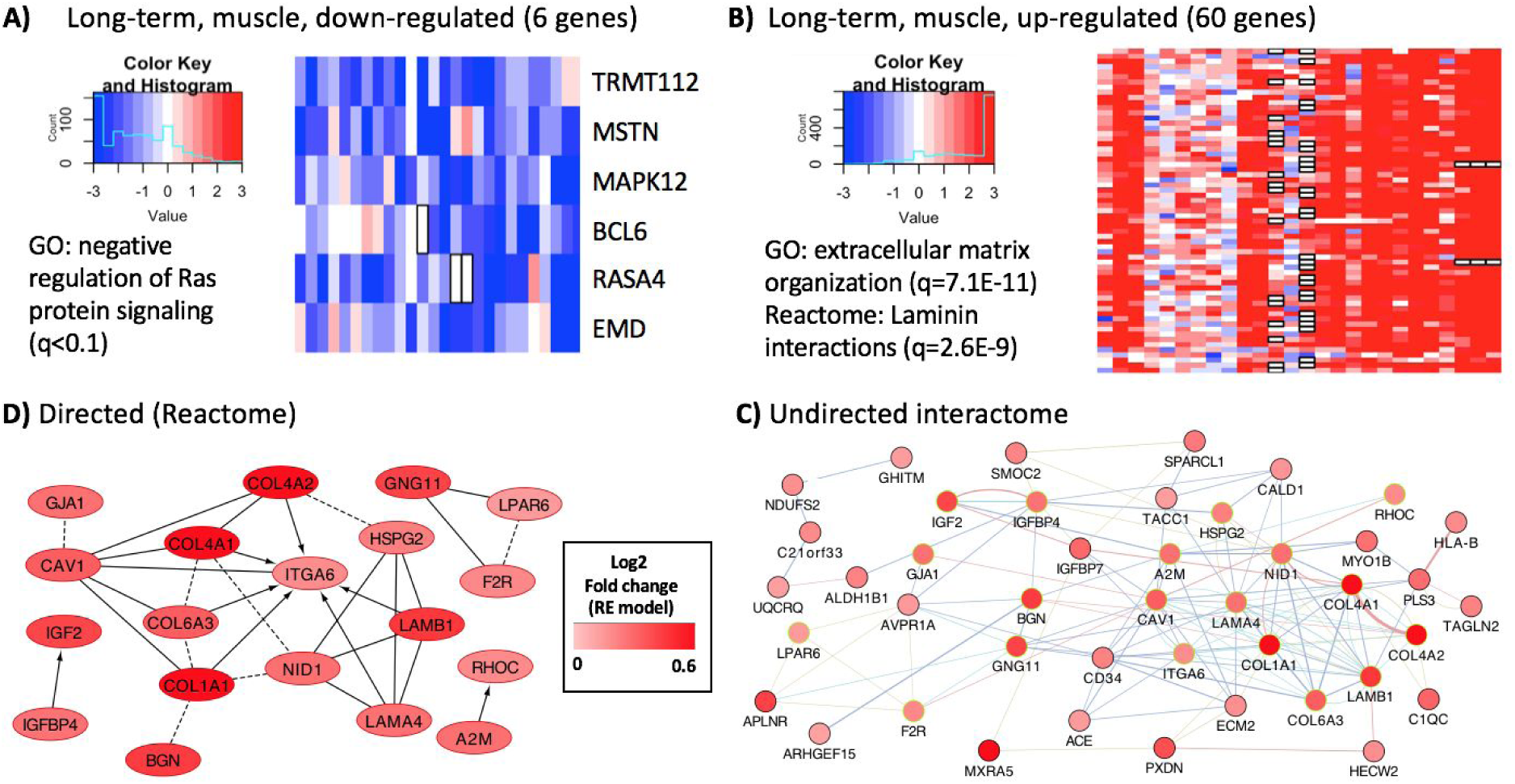
Long-term training muscle response genes with moderator-independent patterns. A) Heatmap of six downregulated genes. B) Heatmap of 60 upregulated genes. C) The main connected component linking most of the upregulated genes to known protein-protein interaction networks. Highlighted nodes are genes with known pathway interactions in Reactome. D) The laminin interactions Reactome pathway covers only a small part of the upregulated gene module. Dashed lines represent co-occurrence in a complex. Directed edges represent regulation.

Although repeated bouts of acute exercise over a longer period of time evidently lead to adaptation, the transcriptional overlap between acute and long-term exercise responses was very low. Only three genes were differentially expressed in response to both acute exercise and long-term training in skeletal muscle; *DDN*, *ID3* and *PLEKHO1*. Dendrin (*DDN*) was upregulated in response to both an acute bout of exercise and long-term training, with a significantly greater induction in response to resistance exercise (see Supplementary Table 2). Its function in muscle is largely unknown, but differential expression has been found in rodent cerebellum and spinal cord with exercise (Caetano-Anollés et al., 2016; Perreau et al., 2005). *ID3* is a transcriptional regulator that inhibits skeletal muscle differentiation and promotes muscle precursor cells proliferation (Baas et al., 2012; Liu et al., 2002; Wu and Lim, 2005). Its expression is negatively correlated with skeletal muscle power in older individuals (Buford et al., 2011) and we also observe a small, but significant association with age. *ID3* is induced by resistance exercise, and to a greater extent in females (Liu et al., 2010). *PLEKHO1* expression increased in response to both acute and long-term resistance exercise interventions. *PLEKHO1* has been found to be a regulator of myoblast elongation and fusion through its interaction with Arp2/3 (Baas et al., 2012).

### Exercise induces more pronounced inflammatory response in older individuals

The sample size, and heterogeneity of the cohorts regarding sex and age allowed us to interrogate potential moderator-specific transcriptional alterations in response to long-term training in skeletal muscle. Differential expression of 22 genes was significantly associated with age (Figure 5A). Particularly, we found that older individuals showed greater induction of interferon-induced inflammatory genes, including *IFITM1*, *GBP1*, *IFI44* and *IFI44L*. Genes associated with degradation of extracellular matrix (*CTSK*, *CD44* and *COL15A1*) were also training-induced specifically in older cohorts.

**Figure 5.**
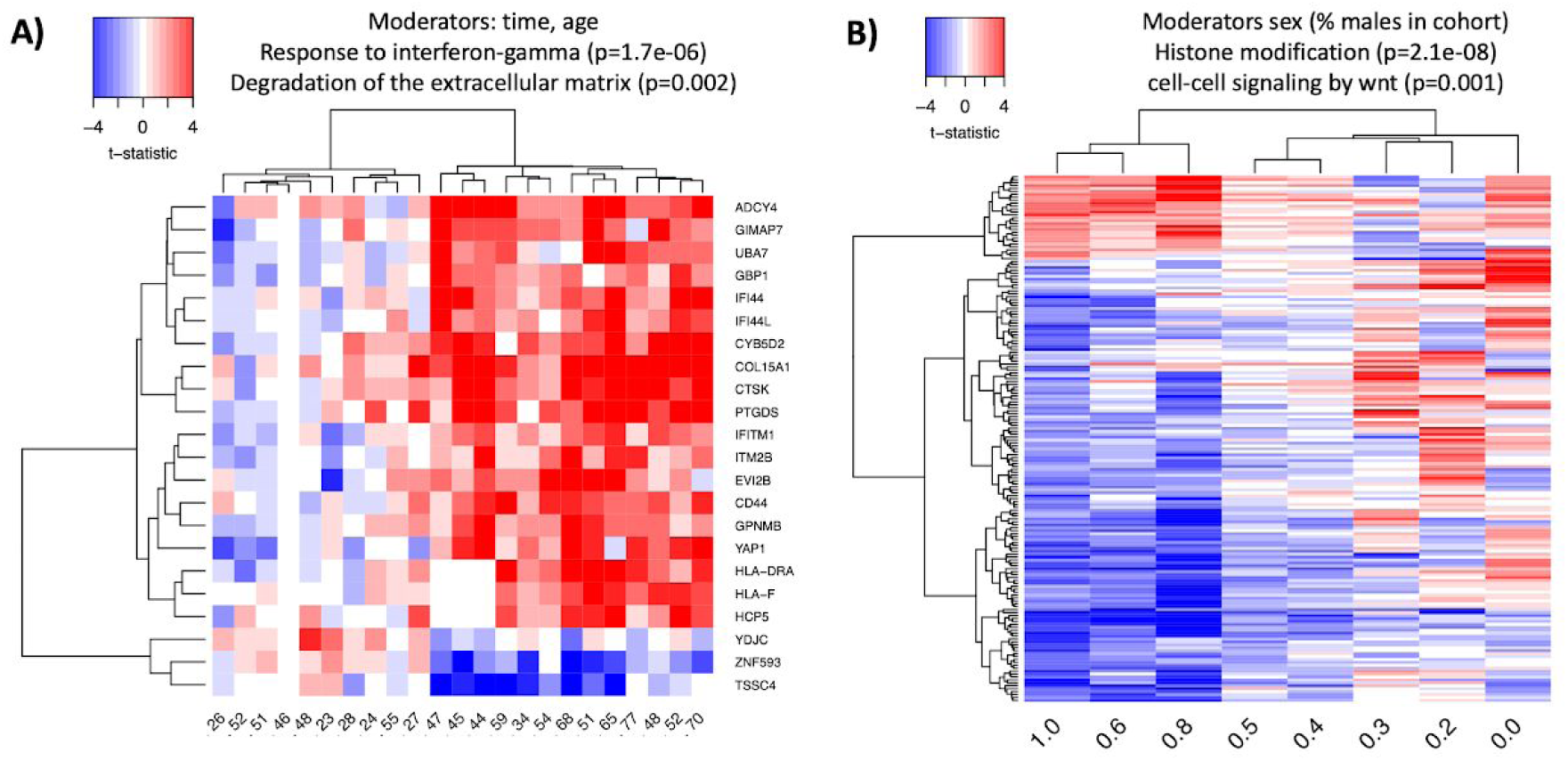
Covariate-specific transcriptional changes in response to long-term training in human skeletal muscle. A) Heatmap of the 22 genes that were differentially regulated based on time and age. B) Heatmap of the 179 genes that were associated with sex distribution within a cohort. In both maps, each cell is the average t-statistic of a gene across a set of studies.

With the 276 males and 197 females that were included in the long-term skeletal muscle meta-analysis (18 subjects were also included without sex-information), we were able to identify multiple genes with sex-specific regulation (Figure 5B). The top functional ontology was histone modification (Supplementary Table 7), with 20 differentially regulated genes. Among them were the methyl-CpG binding protein MECP2, and histone deacetylase 3 (HDAC3), which are important transcriptional regulators in skeletal muscle (Song et al., 2019). An example of a novel exercise-regulated gene is *MTMR3*, a lipid phosphatase that decreased more in male-versus female-predominant cohorts. The same was observed for *AKAP2*, an anchoring protein that regulates PKA signaling. Beta-2-microglobulin (*B2M*), a component of the major histocompatibility complex, increased more in cohorts with more females. Although the decrease in *MTMR3* was small, it was consistent and novel, which made us select this gene for validation. No significant change was observed in the validation cohort, however there was a trend towards a decrease at 6h post exercise (Mean ± SE Pre 0.09±0.01, 6h 0.07±0.007, *P*=0.08), emphasizing the need for the additional power provided by the meta-analysis to discover more subtle gene expression changes.

**Figure 7.**
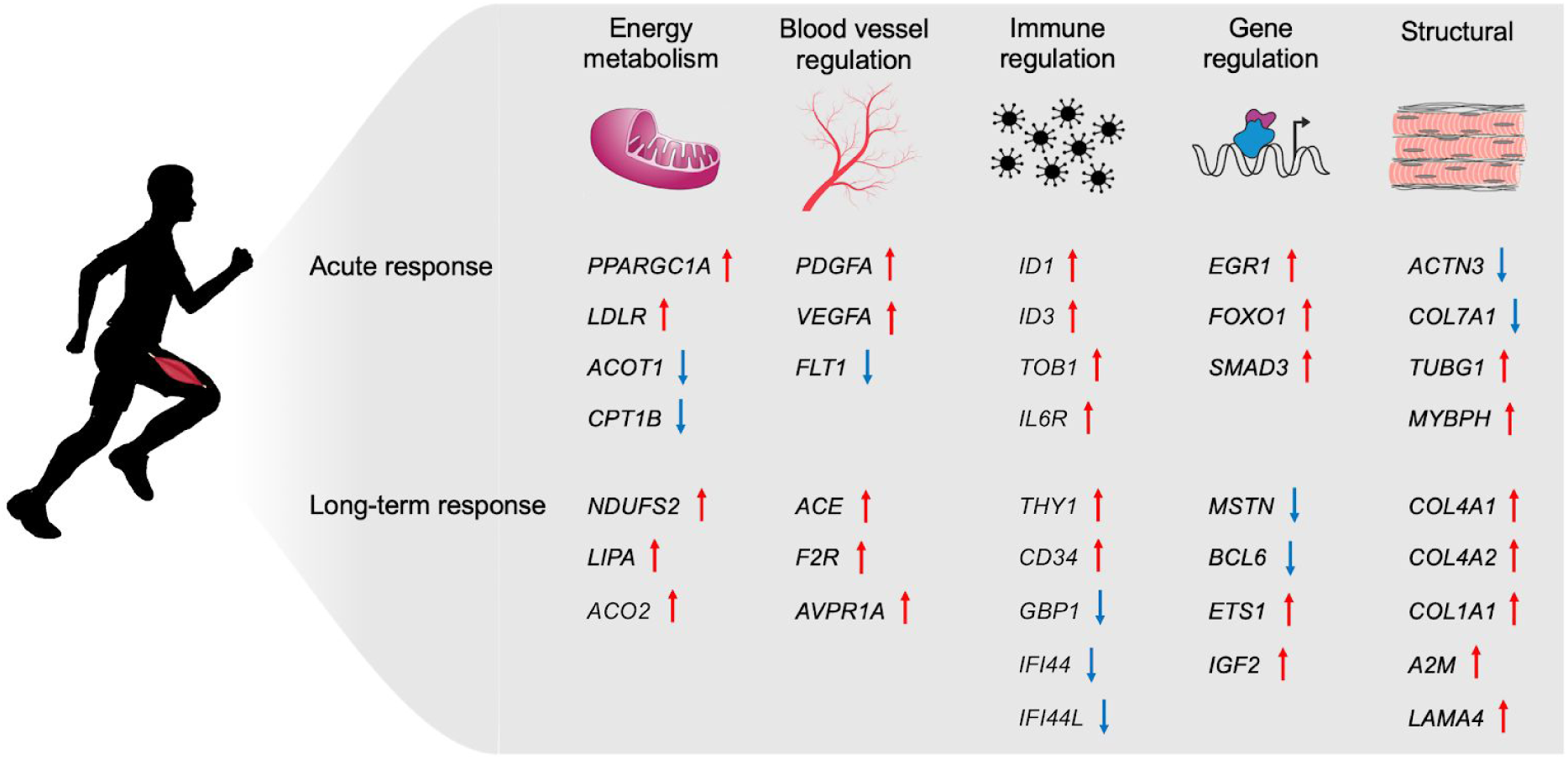
Overview of selected gene expression changes associated with key known adaptation mechanisms in skeletal muscle. Arrows indicate direction of change based on the base model from the meta-analysis.

### Transcriptional response to an acute exercise bout in blood is moderated by age

Almost 400 genes were differentially expressed in blood after acute exercise. Functionally, these genes were associated with exocytosis, activation of leukocytes, and activation of neutrophils (Supplementary Table 7). Interestingly, we found that many of the genes were regulated in the 2-5h window post exercise. However, it is important to note that there were fewer studies covering this time window as compared to 0-1h (4 vs. 13), and there was a bias towards older ages in the >20h time window (Supplementary Table 1, Supplementary Figure 8).

Analyzing the genes associated with time response (Supplementary Figure 7), we found genes associated with RNA processing and translational termination that were downregulated 2-5h after exercise. Many genes coding for ribosomal proteins belonged to this category (e.g. *RPL17*, *RPS17*, *RPS25* and *RPL39*). Mitochondrial ribosome genes (*MRPL18* and *MRPL58*) and *LONP1*, which degrades misfolded proteins within the mitochondrial matrix, were similarly reduced.

Interestingly, the majority (260 out of the total 338) of the differentially expressed genes were moderated by age or a combination of time and age (Figure 6). Genes that interact with voltage-gated Ca-channels (*REM2, CACNB4* and *NOS1AP*) were induced after acute exercise specifically in older cohorts (Figure 6A). Three genes associated with protein destabilization showed a similar pattern (*ID1*, *SIRT6* and *BMP2*). Sirtuin 6 (*SIRT6*) is known to be induced by acute exercise in blood (Chilton et al., 2014) and its expression is associated with aging, an effect that is attenuated with training in rodent skeletal muscle (Koltai et al., 2010).

**Figure 6.**
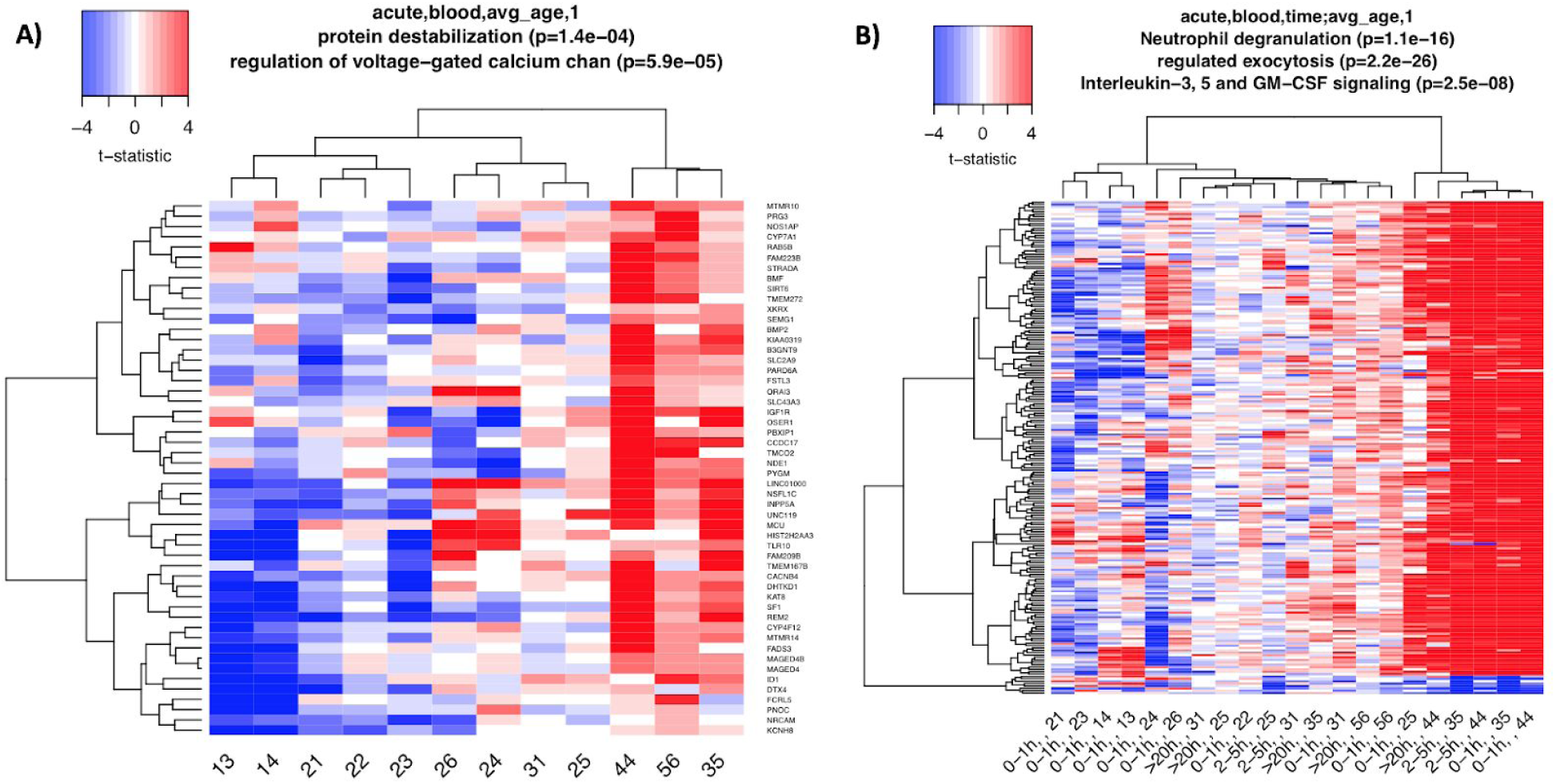
Covariate-specific transcriptional changes in response to acute training in human blood. A) Heatmap of the 52 genes that were differentially regulated based on age. B) Heatmap of the 208 genes that were associated with time and age. In both maps, each cell is the average t-statistic of a gene across a set of studies.

Genes associated with signaling through exocytosis and endocytosis increased more with higher average age in a cohort, as well as with later time windows post exercise (Figure 6B). Myeloid and leukocyte cell activation showed a similar pattern and many genes were common between these ontologies. Examples of common genes were the acute phase anti-inflammatory proteins *ORM1* and *ORM2*, that have been previously shown to be induced by endurance exercise in human blood (Abbasi et al., 2014). Receptors for the anti-inflammatory cytokine IL4 and the growth factor IGF2 were also common between the ontologies.

## Discussion

We took a meta-analysis approach to build a transcriptional landscape of acute and long-term exercise adaptation in human skeletal muscle and blood. We identified co-expressed gene clusters that show differential temporal response patterns after acute exercise and indicate information propagation through the gene network. We predicted upstream regulators of the temporal gene clusters and built a more comprehensive response interactome for long-term training in skeletal muscle. We identified sex- and age-associated differential expression, where long-term training induced a more prominent inflammatory response in older individuals. The overlap between the response to acute and chronic exercise was very low, indicating that more data are required to delineate the interactions between acute and long-term training effects. In blood, the majority of the acute transcriptional responses were associated with age. Very few genes were differentially expressed with long-term training in blood.

In skeletal muscle, the acute response showed changes related to immune regulation, structural changes and energy metabolism. The long-term response was dominated by structural changes in the muscle. Interestingly, genes driving angiogenesis were induced acutely, whereas genes related to blood pressure regulation were induced in response to long-term training. Mitochondrial factors, including ATPases, electron transport chain components and TCA cycle enzymes were not found to be significant. However, this may be in part due to our stringent statistical analysis that focused on the top differential expression genes. A summary of selected skeletal muscle results are shown in Figure 7. In blood, the results were not as coherent as in skeletal muscle. Most acute studies covered the early time window (0-1h), while most genes were found to be differentially expressed 2-5h post exercise. We were unable to identify any public transcriptome datasets for acute resistance exercise interventions in blood, where future research efforts are needed.

At baseline, there are large sex differences in the human skeletal muscle transcriptome (Lindholm et al., 2014b; Maher et al., 2009). However, there are only a few studies that have reported sex-differences in transcriptional regulation with exercise, and these have only looked at specific genes with qRT-PCR (Smith et al., 2009; Vissing et al., 2008). Our meta-analysis substantially expands the catalog of sex-associated exercise responses, and offers multiple candidates for further research.

In skeletal muscle, we identified 179 genes that were associated with long-term training response and sex. They were functionally associated with histone modifications, which has been shown to be a key transcriptional regulation mechanism in response to endurance training in human skeletal muscle (McGee et al., 2009).

Studies investigating age-associated differential expression following exercise at a global level are also limited in numbers (Melov et al., 2007; Robinson et al., 2017; Su et al., 2015). (Robinson et al., 2017) identified age-specific responses to high intensity interval training and resistance training, while (Melov et al., 2007) showed that 6 months of resistance training reversed the age differences in gene expression found at baseline between young and old skeletal muscle. (Su et al., 2015) performed a comprehensive meta-analysis of general age-associated transcriptional changes in skeletal muscle, where some genes were associated with physical performance. Several studies have also investigated specific exercise-induced age-associated expression changes using qRT-PCR (Raue et al., 2007; Short et al., 2005). We observed age-associated changes in response to acute exercise in blood (52 genes) and with long-term training in skeletal muscle (22 genes). Immune response and inflammatory genes were induced to a greater extent in skeletal muscle in cohorts with older individuals. It is not clear how a greater inflammatory response at the transcriptional level affects the adaptation in older individuals, as inflammation is an essential part of exercise adaptation, and may also affect the degree of training-induced muscle damage.

Meta-analysis methods have some limitations that should be considered, especially when studies are highly heterogeneous, as we observed in our case. First, random effects meta-analysis does not directly explain variability. Second, meta-regression is often exploratory in nature. Third, when the number of studies is moderate or small (e.g., 15 or less) and their heterogeneity is high then p-values are often biased downwards and small studies may be assigned too much weight (Hippel and von Hippel, 2015; Serghiou and Goodman, 2019). This observation was also noted previously in the gene expression case, where selecting genes based on the p-value of the meta-analysis resulted in thousands of genes and poor replicability even after false discovery rate adjustment (Sweeney et al., 2017). Finally, meta-analysis can suffer from low power when the heterogeneity is high or when studies are sampled in an unbalanced way across their moderators (Hempel et al., 2013; Rubio-Aparicio et al., 2017).

We mitigated these issues by introducing a model selection pipeline that fits the moderator set for each gene to better explain the variability in the data. Moreover, we do not rely on p-values alone to select genes: we require that their models will have both a high AICc difference and a high effect size (fold change). As most moderator pairs are not correlated (Supplementary Figure 1), we expect that the model selection process correctly detected the relevant moderators in most cases. However, in the acute bout blood data we observed some biases and correlated moderators: fewer studies covered the intermediate time window, and there was a bias towards older ages in the >20h time window. Nevertheless, the detected genes in all of our analyses had high cross-study replicability when evaluated using empirical Bayes algorithms (Amar et al., 2017; Heller et al., 2014). Thus, while some specific models may not be accurate, the selected genes are consistently associated with response to exercise. This is corroborated by the high concordance between the meta-analysis findings and the results from the validation cohort.

Understanding the underlying exercise response mechanisms holds immense potential for human health and medicine. This meta-analysis of the transcriptomic response to exercise reveals differential expression trajectories in skeletal muscle and blood. In skeletal muscle, we find co-expressed gene clusters that show differential temporal response patterns after acute exercise with information propagation through the gene network. We identify multiple age- and sex-associated differentially expressed genes, which provides important knowledge for improving exercise prescription for precision health. Overall, this meta-analysis deepens our understanding of the transcriptional responses to exercise and provides a powerful, free and easily accessible public resource for future research efforts in exercise physiology and medicine.

## Materials and Methods

### Data collection

We inspected and annotated publicly available human exercise omics data available in the Gene Expression Omnibus (Edgar et al., 2002) (GEO, search date 2/22/2019). The datasets were annotated by looking at their metadata taken from GEO or Recount (Frazee et al., 2011), by extracting additional information from the publications (including supplementary data), and (whenever required) by personal communication with the authors. Only transcriptomics data had a reasonable number of studies to allow a meta-analysis of differential abundance. For these meta-analyses, we collected human blood and skeletal muscle gene expression datasets that had pre- and post-exercise samples for most of their subjects. Non-transcriptomics data, adipose tissue datasets, and datasets without multiple time points per subject were excluded from the analysis, even though some are available in the database (Supplementary Table 1). Each dataset was partitioned into cohorts based on tissue, study arm, and training modality. This was performed separately for acute bout datasets and long-term datasets (some studies had both). However, note that datasets GSE59088, GSE28998, GSE28392, and GSE106865 had both acute and long-term cohorts. The included long-term studies ranged from six weeks to nine months of training.

After our manual preprocessing above, 413 samples had missing sex information. To address this we took a machine learning approach to impute the missing sex values. We trained a linear SVM classifier for sex using the expression levels from 469 Y chromosome genes that were covered by most of the platforms in our data. Studies from platforms GPL571 and GPL16686 (long-term cohorts 26, 31, and 35 and the acute cohort 37) were excluded due to low overlap with the other platforms, see Supplementary Table 1. The remaining data contained a training set of 1622 samples with known sex information (1082 males). Internal leave-study-out cross validation had very high performance with 0.99 area under the ROC curve and 0.96 area under the precision-recall curve. The predictions of final model (trained using all labeled data) on the 413 samples with the missing sex information (225 of which were predicted to be males) were used for the meta-analysis.

### Preprocessing and moderators

For each gene in each cohort we computed the mean and standard deviation of the log fold-change between the post-exercise time points and the baseline, illustrated in Figure 1. We also kept the p-value of the pairwise test for each time point (i.e., vs. the pre-exercise samples). The result is a matrix of genes over the summary statistics (y – the mean log2 fold change, v – the log2 fold change variance, p – the paired t-test p-value).

For each cohort the following moderators (covariates) were collected: experiment type (long-term training program vs. single acute bout), tissue (blood vs. muscle), training modality (endurance, untrained, or resistance), sex measured as the proportion of males in the cohort, average age, and time (measured in hours for acute bouts and days for long-term training). The cohorts did not cover all combinations of the moderators. Moreover, the low sample sizes, limited number of cohorts, and the observed heterogeneity, all posed challenges that may create inflation in false positives. Below we explain the different steps we took to select cohorts and moderators for the analysis where the goal was to both select relatively homogeneous cohorts and allow exploration of some moderators that were reasonably covered.

Exercise vs. untrained cohorts: we first observed that most exercise studies did not monitor the gene expression change in untrained controls. We kept these cohorts separately (5 for acute, muscle; 2 for acute, blood, and 2 for long-term, muscle, <30 subjects in each case) and used them to validate the selected genes from the meta-analyses (Figure 2E).

Next, for the exercise cohorts we examined the moderators in each case and decided to include the following moderators in each analysis:

1. Acute, blood (17 cohorts in total): time, sex, and age. Time was partitioned into early response (≤ 1 hour post exercise), intermediate (2-6 hours), and late (≥ 24 hours). Training was excluded because no cohort had resistance training.
2. Acute, muscle (15 cohorts): time, training, and age. Time was partitioned into windows as in the acute blood analysis above. Sex was excluded as only four cohort had females.
3. Long-term, muscle (29 cohorts): all moderators. Time was binned into < 150 days, and >150 days.
4. Long-term, blood: No moderators, this analysis had only three cohorts.

### Meta-analysis of a single gene

For meta-analysis (or meta-regression) we excluded genes that had a missing value in 25% or more. This still left more than 18,000 genes in each of the analyses (acute muscle - 18,374, acute blood - 18,621, long-term muscle - 18,685, long-term blood - 19,683).

#### Random-effects meta-analysis

We used the R metafor package to perform meta-analysis and meta-regression for each gene (Viechtbauer, 2010). In both cases, we model the fold change of a gene. Assume we are given *y_i_*, *i* ∈ 1, …, *k* estimated effects of a gene, where *i* denotes a specific post-exercise time point in a cohort (nested within studies). Let *y_i_* be the log fold change with variance *v_i_*. In a standard random effects meta-analysis we assume that *y_i_* = μ + *u_i_* + ε*_i_*, where *y_i_*, is the observed effect, *u_i_* is the true (unknown) effect *u_i_* ∼ *N* (0, τ ^2^), and ε*_i_* is the sampling error term, ε*_i_* ∼ *N* (0, *v_i_*), which are assumed to be independent. In this analysis we were interested in both the fold change of the gene across all cohorts and time points (μ), and the total amount of true heterogeneity among the true effects (τ ^2^). True effects heterogeneity is important in meta-analysis as it serves as a measure of consistency. A standard interpretable statistic of consistency is *I*^2^ : the ratio of true heterogeneity to total observed variation (Higgins et al., 2003).

As explained above, in a standard meta-analysis the random effects are assumed to be drawn independently from *N* (0, τ ^2^). However, we can incorporate a specific structure of these effects. Following the notation in the metafor package we use a nested model structure, denoted as *inner|outer.* Observations with the same *outer* value follow some structure based on their *inner* value. For example, if *inner* is the cohort and *outer* is the study, we can set the random effects to have a block-wise structure: observations from different studies will be independent, but observations from cohorts from the same studies will be dependent using a defined variance-covariance structure. In our analyses we tested either the standard error form or a cohort|study random effects model with a general structure (“CS”).

Random-effects meta-analysis was used both as a base model to evaluate the effect and heterogeneity in the exercise meta-analyses (1-4 above), and to evaluate the fold change of a gene in the untrained controls. The goal of analyzing the untrained controls was to evaluate genes that are differential but not because of the exercise (e.g., due to the process of taking a muscle biopsy).

### Meta-regression

We used mixed-effects meta-regression to extend the analysis to include a set of effect moderators (i.e., covariates). In this case, assuming that there is a large unexplained heterogeneity among the true effects, we model the fold change of a gene using:

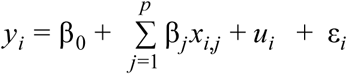

where *x_i_*_,*j*_ denotes the value of the j-th moderator variable for the i-th effect, *u_i_* represents a random effect, whose variance represents the residual heterogeneity among the true effects. In other words, it is the variability among the true effects not captured by the moderators in the model. The different tested distribution structure of *u_i_* are the same as in the meta-analysis explanation above.

### Model selection

Whenever meta-regression was considered, we tested all possible combinations of a subset of the moderators (including no moderators and all moderators as options). This was combined with testing two different random effects structures as discussed above. For each model we computed the AICc score, which is more suitable for our case with a limited number of studies as compared to other alternatives (Burnham and Anderson, 2002; Cavanaugh, 1997). We then ranked all models by their AICc and excluded models that were not among the top two or were not base models (i.e., models without moderators).

### Gene selection process

To reduce the number of model selection analyses, we first excluded genes that had no or only a single p-value < 0.05 across all cohorts. For the remaining genes we computed the base models and performed the model selection analysis.

A model was considered a significant improvement over a base model (and thus selected) only if Δ*AICc* > 5, *p* < 0.001, ∃β*_j_*; |β*_j_* | > μ_0_, where *p* is the p-value of the model (including all moderators), and μ_0_ is a threshold for absolute effect size. We used μ_0_ >0.25 for acute studies, and μ_0_ >0.1 for long-term.

In case that no improvement over the base model was identified, the base model was selected only if it had *I*^2^ < 50%, *p* < 0.001, |μ| > μ_0_, where *p* is the p-value of the estimated effect and μ_0_ is defined as above.

Note that using both thresholds for the model significance and the effect size is recommended as it tends to reduce false positives and highlight reproducible results (Sweeney et al., 2017).

### Replication analysis

Replication analysis is a different type of integrative analysis whose goal is to directly estimate replication of observations by inspecting their p-values alone (Heller et al., 2014). Such analysis addresses challenges that are typically not covered by standard meta-analysis: (1) latent correlation structure among the observations, (2) quantifying replication directly, and (3) quantifying the FDR either of single genes (called local FDR) or of a gene set (with respect for their replication).

We used the SCREEN algorithm (Amar et al., 2017) to evaluate the Bayes local FDR of genes relative to different replication thresholds. SCREEN takes as input a matrix of p-values or z-scores, and a range 1, …, *k*. For each gene and a replication level *k* ∈ 1, …, *k*′ SCREEN estimates an upper bound to the local FDR of a gene defined as:

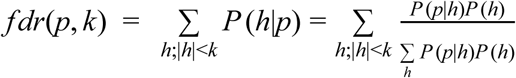

where *p* is the set of p-values of the analyzed gene, *h* is a binary vector of size n, *P* (*h*) is the prior distribution of *h* (estimated from the data).

### Gene clustering

In each of the four analyses (e.g., acute muscle) we took all models that had the same covariates and clustered their t-statistic matrices using *K-means* (Hartigan and Wong, 1979). In each such analysis we determined the number of clusters using the elbow method: we plot the within-cluster sum of squares as a function of the number of clusters (testing 1-15) taking the last point that had at least 60% improvement from the previous one as the number of clusters. For the time course model detected in acute muscle we also used DREM for clustering and network inference for transcription factors (TFs) (Schulz et al., 2012). Here, we used DREM with the ENCODE TF-target network (Harrow et al., 2012) and a low penalty for adding nodes (5 instead of the 40) as the default value did not result in a clustering of the time courses.

### Functional analysis and network visualization

GO enrichment analysis was performed using topGO (Alexa and Rahnenfuhrer, 2010). Reactome enrichment analysis was performed using ReactomePA (Yu and He, 2016). Network analysis for selected gene sets was completed using GeneMANIA (Warde-Farley et al., 2010) in Cytoscape (Shannon et al., 2003; Smoot et al., 2011).

### Validation cohort and qRT-PCR

A separate acute endurance exercise cohort of 16 individuals, 8 males and 8 females, was included to validate some of the meta-analysis findings. The study, which was approved by the Regional Ethical Review Board in Stockholm, Sweden, has been described elsewhere (Gidlund et al., 2015). In brief, the subjects performed 60 min of endurance exercise on a cycle ergometer at a workload corresponding to 70% of their peak VO_2_. Skeletal muscle biopsies from *vastus lateralis* were obtained before exercise, and at 2h and 6h after the exercise bout. Total RNA was extracted using the acid phenol method (Chomczynski and Sacchi, 1987) and 1ug RNA was reverse transcribed using Superscript reverse transcriptase (Life Technologies) and random hexamer primers (Roche Diagnostics). Quantitative real-time PCR was performed with primers for SMAD3, NR4A1, SCN2B, HES1, ID1, SLC25A25, MTMR3 (Sigma-Aldrich, primer sequences are listed in **STable 9**) and with Glyceraldehyde-3-phosphate dehydrogenase (GAPDH, 4352934E, Applied Biosystems) as an endogenous control gene. Expression level was calculated using the ΔC_T_ method and statistical analysis performed using a mixed effects model followed by Dunnett’s multiple comparisons test.

### Data and code availability

All data and associated code are available here:
https://github.com/AshleyLab/motrpac_public_data_analysis

## Acknowledgements

This research was supported by the NIH Common Fund (award number U24OD026629) and the Knut and Alice Wallenberg Foundation (M.E.L).

## Author contributions

Conception and design: DA, MEL, MTW, MR, EAA. Analysis and interpretation: DA, MEL, MTW, EAA. Validation experiments: JN. Manuscript draft: DA, MEL. All authors contributed to the research of the published work, and have read and approved the final manuscript. The authors declare there is no conflict of interest.

## Supplementary Figures

**Supplementary Figure 1.**
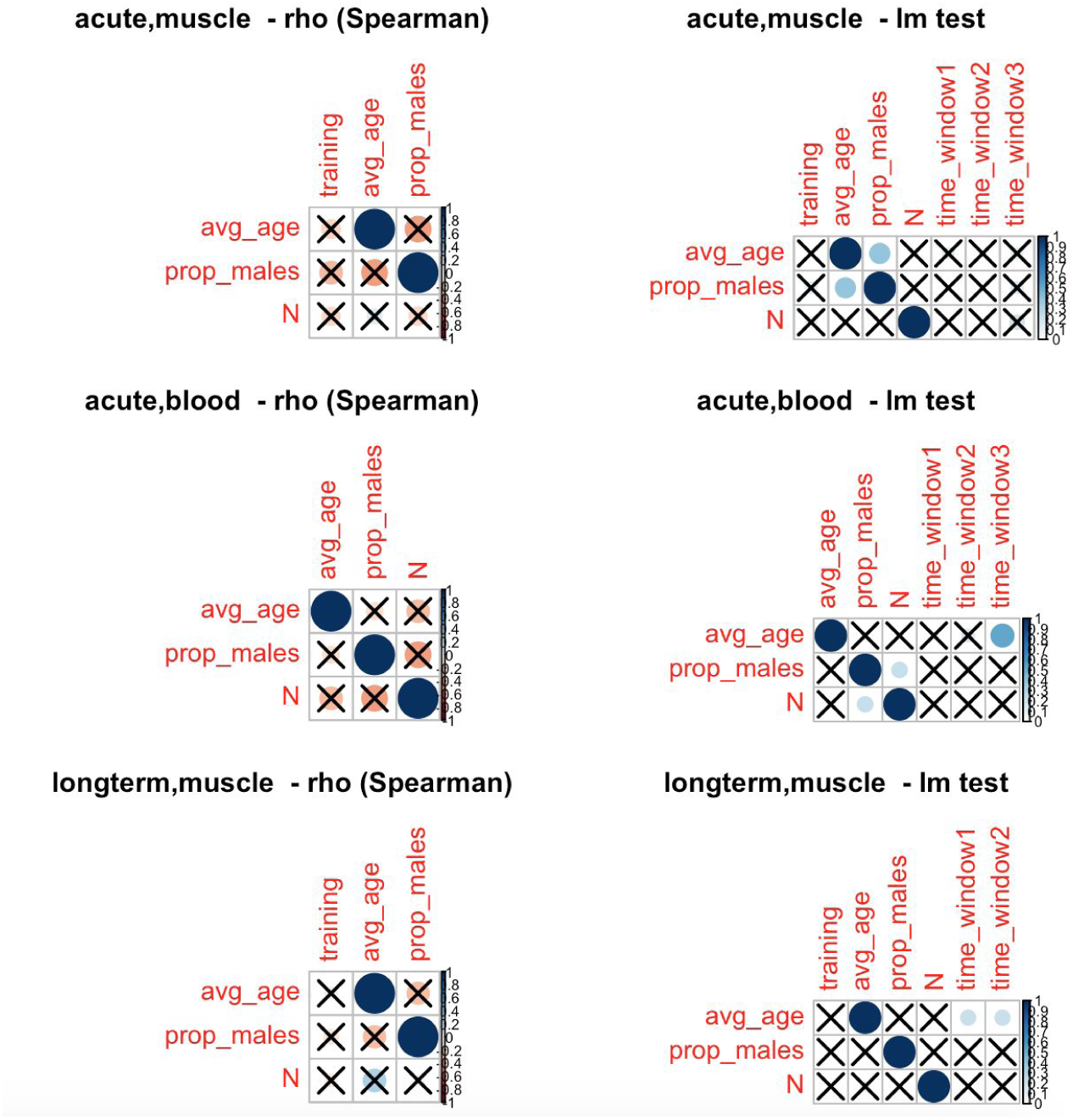
Spearman correlations of moderators (covariates) across the cohorts in our four meta-analyses. N is the number of subjects. For time, acute studies were binned into three windows (1:0-1h, 2: 2-6h; 3:20h+). Long-term studies were binned into two time windows (up to 150 days or greater). X indicates that the correlation between a pair of moderators is not significant.

**Supplementary Figure 2.**
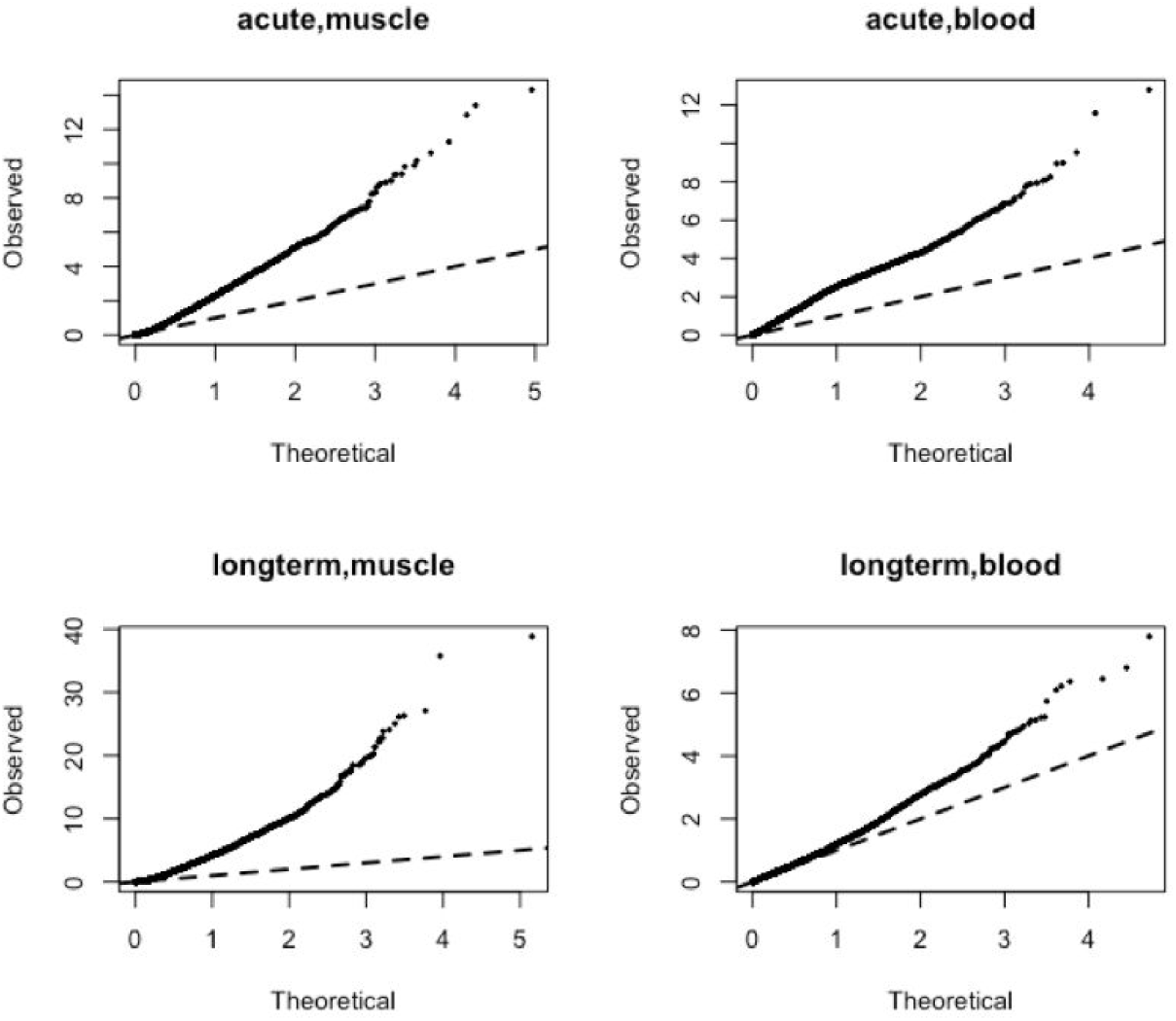
QQ-plot of the random-effects meta-analysis. All cases had >10,000 genes and the observed p-value distribution was inflated with low p-values.

**Supplementary Figure 3.**
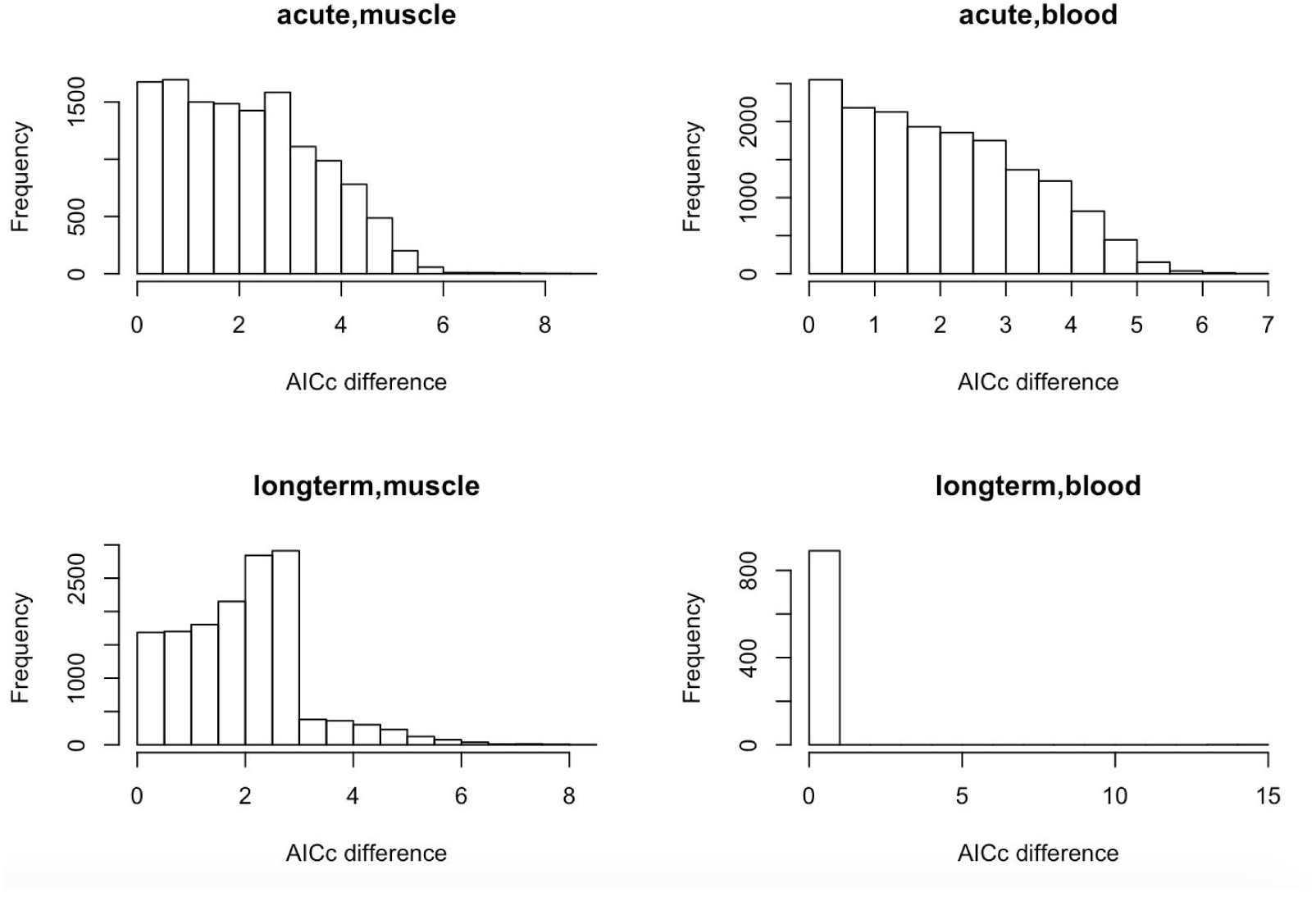
Histograms of AICc scores of the model selection process. For each gene we computed the difference between the base model (random effects, no moderators) and the top model. Each histogram shows the distribution of the AICc differences.

**Supplementary Figure 4.**
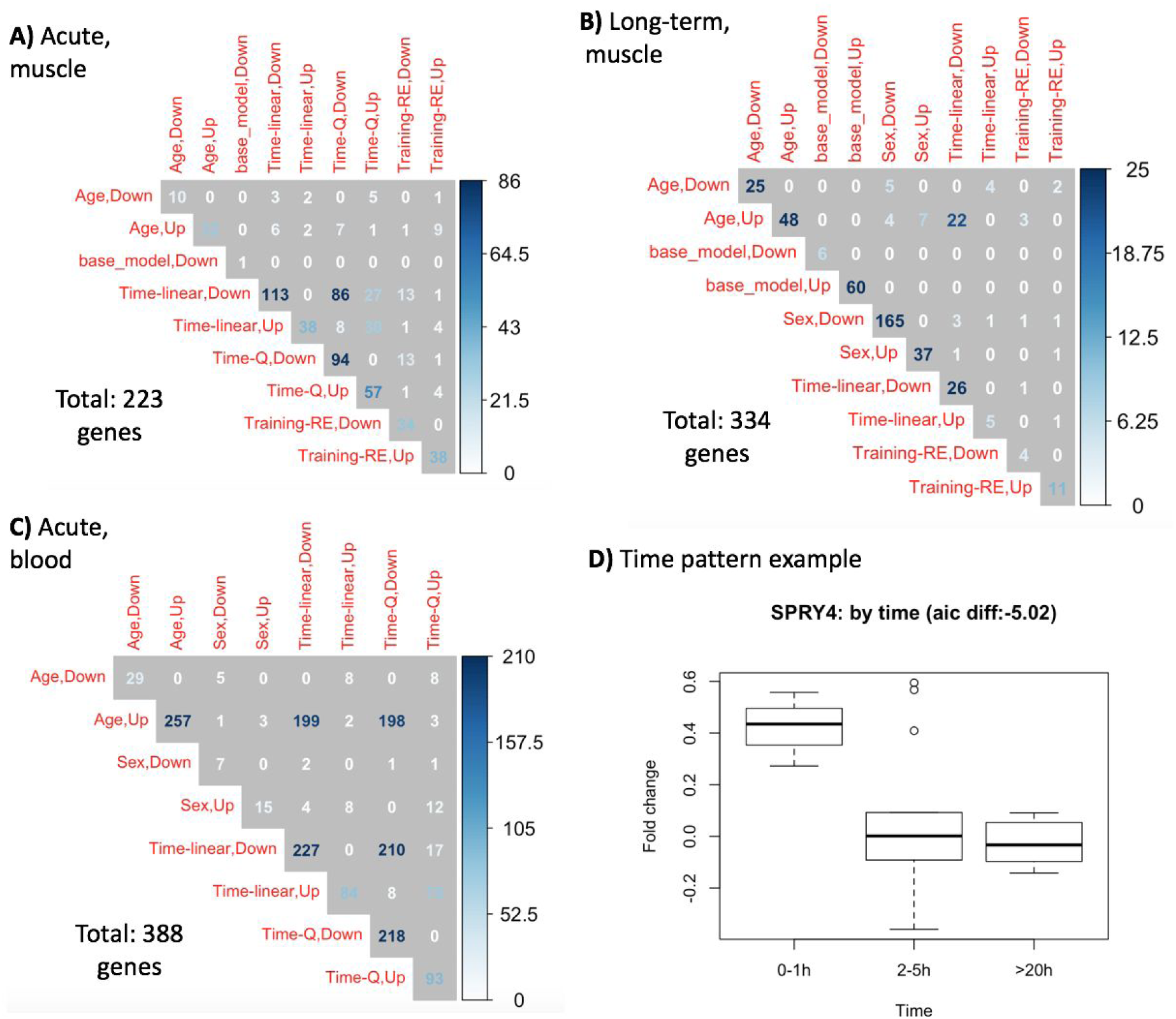
Selected models and their moderators (covariates). A-C) the matrices show the co-occurrence of covariates in selected models. Each number represents the number of genes whose models had the two moderators. The diagonal shows the total number of models for each moderator. Sex: the proportion of males in a study. Time-linear and Time-Q are orthogonal polynomials used to model the time response (e.g., a linear trend will approximate a monotone up- or down-regulation response). Training-RE: a binary covariate specifying if the cohort included resistance training. Up/Down in the covariate name specifies if the regression coefficient is positive or negative, respectfully. D) SPRY4 in acute muscle cohorts is upregulated only in the first time window. In this case the selected model had time associated features that represent both a linear and a quadratic trend.

**Supplementary Figure 5.**
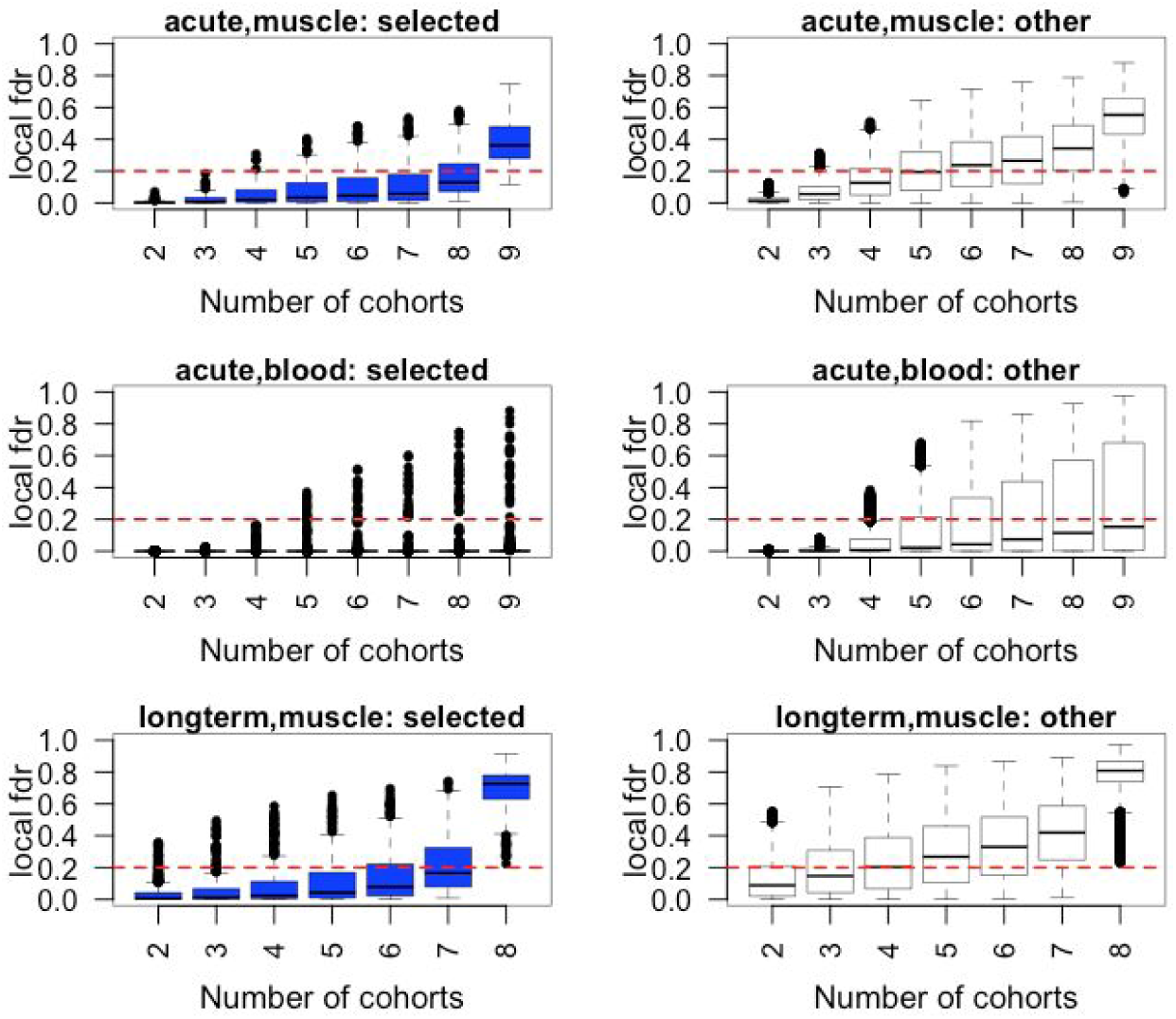
SCREEN analysis illustrates high replicability of the selected genes. Each subplot shows the distribution of the gene local FDR scores as a function of the input replication level. Blue boxplots show the local FDR scores of our selected genes. White boxplots show the local FDR scores of the genes that were tested in the model selection process but were not selected. These genes were a subset of all covered genes in all analysis: they had a significant p-value in at least one time point in at least one cohort.

**Supplementary Figure 6.**
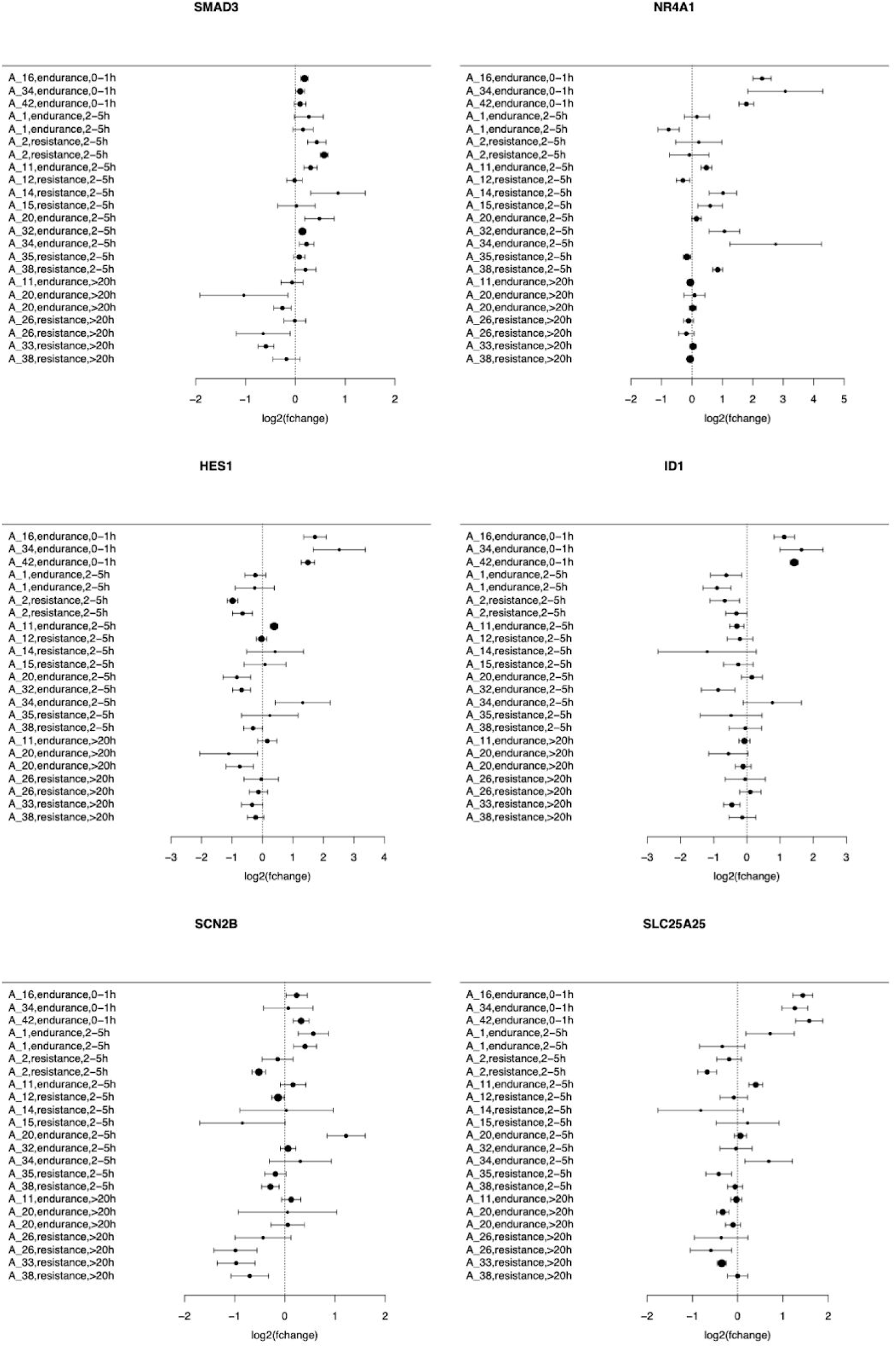
Forest plots for genes selected for validation from the skeletal muscle acute exercise meta-analysis.

**Supplementary Figure 7.**
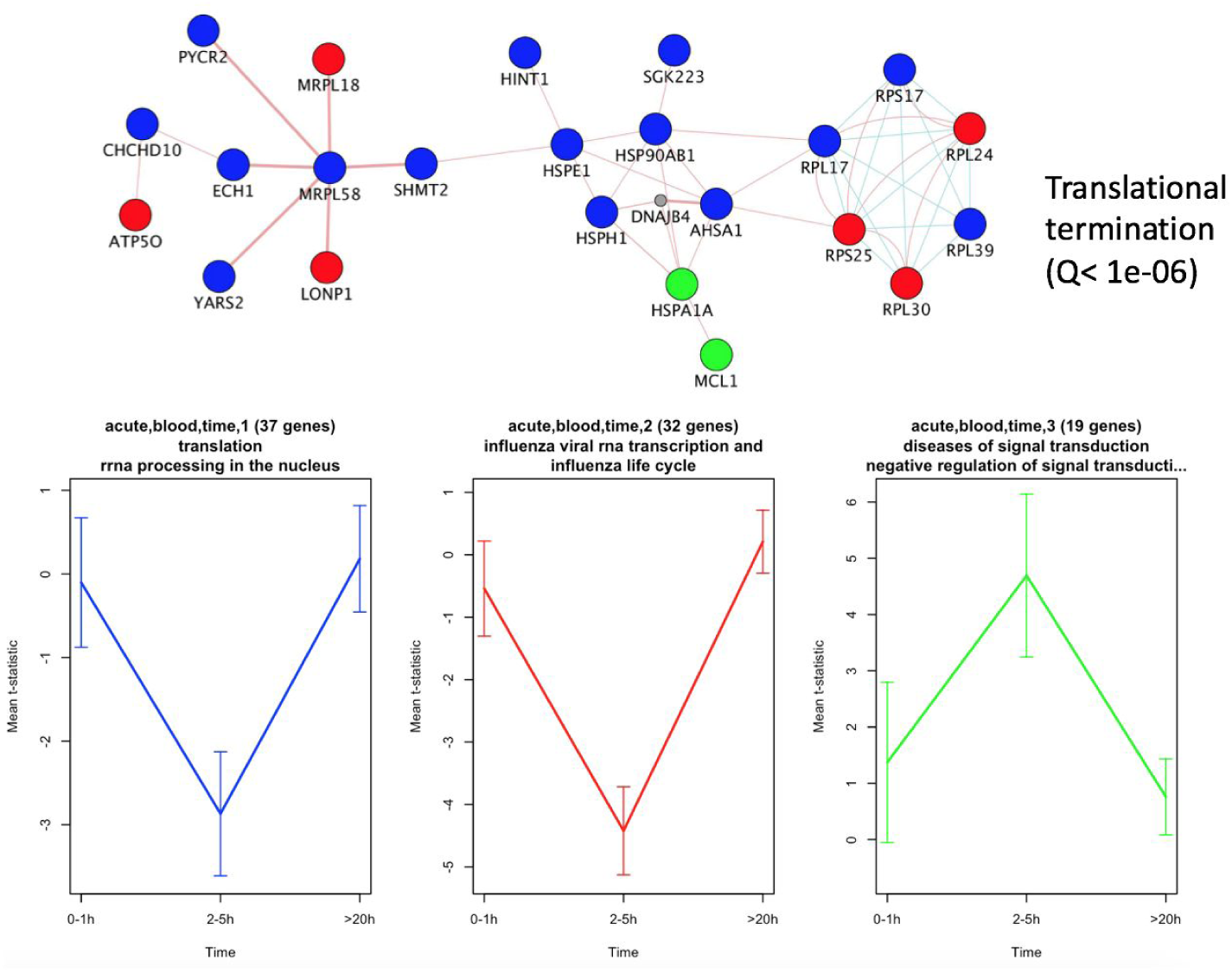
Differential expression patterns in blood after acute exercise. Genes associated with time were partitioned into three groups based on their trajectories (bottom panel). The main connected component of the genes when overlaid on known protein-protein or pathway networks was functionally associated with translational termination (upper panel).

**Supplementary Figure 8.**
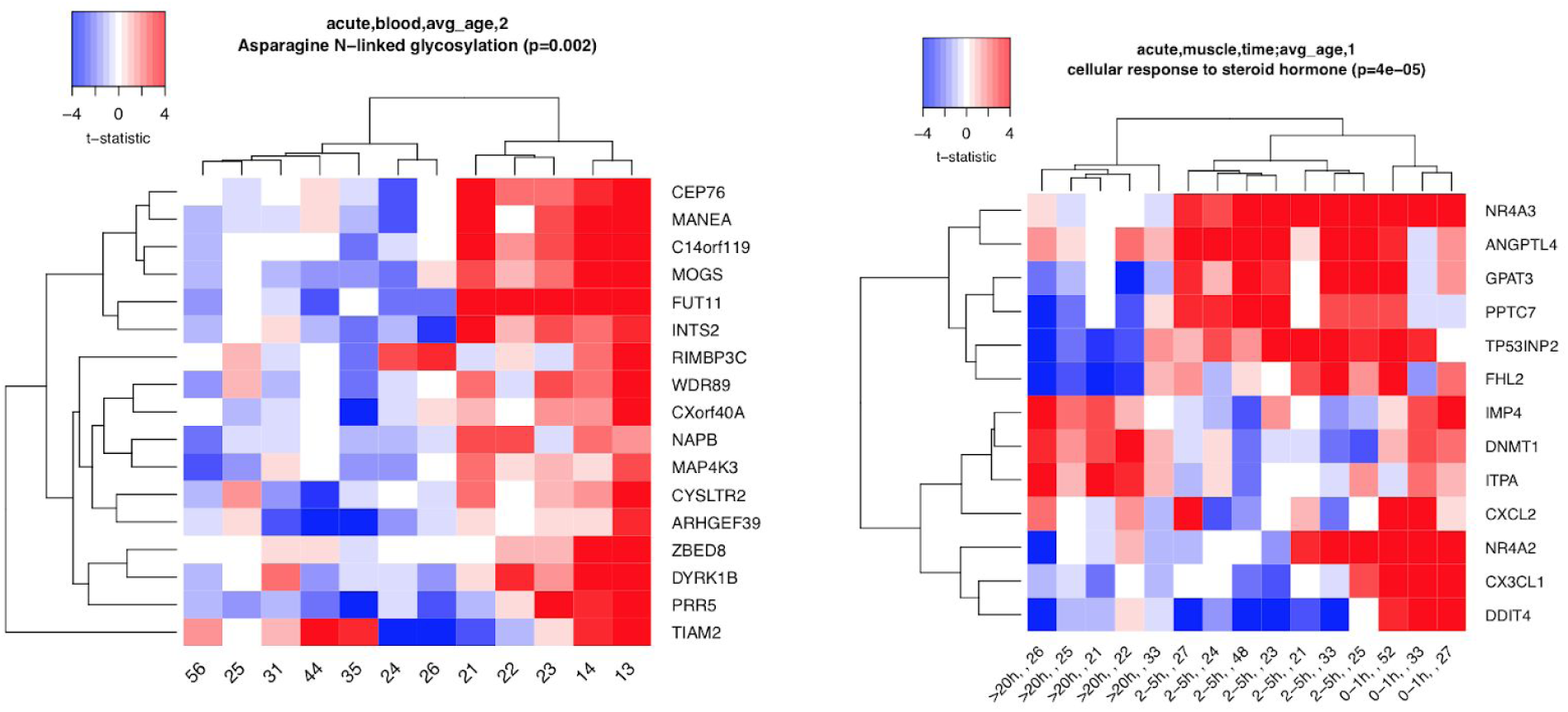
Differentially expressed genes in response to acute exercise in blood. Left panel shows a heatmap of the genes that were significantly associated with average age, and right panel shows genes that were moderated by both time window post exercise and age.

## Supplementary Table Legends

**STable 1.** Cohort metadata and moderators.

**STable 2.** Model selection results. Each row represents a significant (meta-analysis, gene, moderator) triplet. The model’s AICc difference and significance are shown. In addition, the gene clustering results are presented.

**STable 3**. Model selection results in acute muscle: full models. Each row represents a (gene, moderator/intercept) pair. The effect size and p-value are shown for each pair.

**STable 4.** Model selection results in acute blood: full models. Each row represents a (gene, moderator/intercept) pair. The effect size and p-value are shown for each pair.

**STable 5.** Model selection results in long-term muscle: full models. Each row represents a (gene, moderator/intercept) pair. The effect size and p-value are shown for each pair.

**STable 6**. Model selection results in long-term, blood: full models. Each row represents a (gene, moderator/intercept) pair. The effect size and p-value are shown for each pair.

**STable 7.** GO enrichment analysis.

**STable 8.** Reactome pathway enrichment analysis.

**STable 9.** Primer sequences used for qRT-PCR experiments.

